# Antagonistic Regulation of Arabidopsis Leaf Senescence by SCOOP10 and SCOOP12 Peptides via MIK2 Receptor-Like Kinase

**DOI:** 10.1101/2023.10.27.564453

**Authors:** Zhenbiao Zhang, Nora Gigli-Bisceglia, Wei Li, Christa Testerink, Yongfeng Guo

**Affiliations:** Tobacco Research Institute, Chinese Academy of Agricultural Sciences, Qingdao, China; Graduate School of Chinese Academy of Agricultural Sciences, Beijing, China; Laboratory of Plant Physiology, Wageningen University & Research, Wageningen, Netherlands; Plant Stress Resilience, Institute of Environmental Biology, Utrecht University, Utrecht, Netherlands

**Keywords:** Leaf senescence, MIK2, SCOOP10, SCOOP12, Small signaling peptides, Receptor-like kinase

## Abstract

Leaf senescence plays a critical role in a plant’s overall reproductive success due to its involvement in nutrient remobilization and allocation. However, our current understanding of the molecular mechanisms controlling leaf senescence remains limited. In this study, we demonstrate that the receptor-like kinase MALE DISCOVERER 1-INTERACTING RECEPTOR-LIKE KINASE 2 (MIK2) functions as a negative regulator of leaf senescence. We report that the SERINE-RICH ENDOGENOUS PEPTIDES 10 and 12 (SCOOP10 and SCOOP12), previously known to physically interact with MIK2, compete with each other to control MIK2-dependent senescence mechanisms. We observed that increased expression of SCOOP10 or the application of exogenous SCOOP10 peptides accelerated leaf senescence in a MIK2-dependent manner. Conversely, SCOOP12 acted as a suppressor of MIK2-dependent senescence. Thus, the SCOOP10-MIK2 and the SCOOP12-MIK2 modules function antagonistically, allowing for fine-tuning the modulation of the leaf senescence process. Our research sheds light on the complex mechanisms underlying leaf senescence and provides valuable insights into the interplay between receptors, peptides, and the regulation of plant senescence.

## Introduction

Senescence, a crucial biological event in plant development, involves the coordinated breakdown of cellular components to recycle nutrients, aiding adaptation to changing environments. It optimizes nutrient storage and reuse, particularly in leaves, the primary photosynthetic organs. Initiation of leaf senescence prompts various physiological changes, including chlorophyll reduction and macromolecule degradation, enabling nutrient recycling that supports the growth of seeds and fruits^1,2,3^. Efficient senescence is vital for non-perennial plants like wheat and sorghum, as it maximizes nutrient use and prevents yield loss during adverse conditions^4,5^. Understanding the molecular mechanisms behind leaf senescence is critical for improving agricultural productivity, especially in challenging climates. Senescence is a programmed cell death process, and is triggered by endogenous cues and environmental factors, with phytohormones like ethylene, cytokinins, abscisic acid, and jasmonic acid playing significant roles^6^. Recently, peptide hormones have emerged as signaling molecules controlling several aspects of plant developmental processes, including senescence^7,8^. These plant endogenous peptides, typically consisting of fewer than 100 amino acids, are synthesized within the cytosol and subsequently secreted into the apoplast where they are thought to interact with receptor protein kinases located at the plasma membrane^9,10,11^. For example, the CLAVATA3/ EMBRYO-SURROUNDING REGION- RELATED (CLE) peptides family member CLE14 and CLE42 negatively regulate age-dependent leaf senescence^8^. In fact, loss-of-function mutations in CLE14 or CLE42 caused an accelerated senescence while overexpression of *CLE14* or *CLE42* led to a delayed leaf senescence^12,13^. Additionally, the INFLORESCENCE DEFICIENT IN ABSCISSION-LIKE6 (IDL6) peptide has been reported to positively modulate leaf senescence in *Arabidopsis thaliana*. *idl6* mutants display delayed leaf senescence, while overexpression of *IDL6* accelerates leaf senescence^14^. Furthermore, the plant elicitor peptide PEP1, one of the first plant phytocytokines identified to regulate plant immune responses^10,15,16^, has also been shown to function as a leaf senescence modulator in Arabidopsis^17^. SERINE RICH ENDOGENOUS PEPTIDES (SCOOPs) are an emerging class of phytocytokines present specifically in *Brassicaceae*, and have been recently reported to be implicated in controlling plant immunity and development^18–20^. The precursors of SCOOPs, PROSCOOPs^18^ have to undergo active N-terminus proteolysis to generate C-terminus active SCOOP peptides. At least 23 SCOOPs (SCOOP1-23) have been identified in Arabidopsis^20^. Among them, SCOOP12 has been shown to bind to the Leucine-Rich Repeat (LRR) Receptor-Like Kinases (RLK) MALE DISCOVERER 1-INTERACTING RECEPTOR-LIKE KINASE 2 (MIK2) to activate the plant immune response to *Fusarium oxysporum*^19,20^. PROSCOOP10 was recently found to process two encoded regions of PROSCOOP10, resulting in two distinct forms of SCOOP peptides (SCOOP10#1 and SCOOP10#2)^21^. SCOOP10#2 (referred to as SCOOP10 in this article) has also been shown to have dual effects, inducing plant immune responses and inhibiting root elongation^19,20^. MIK2 has been reported to recognize the Fusarium-derived SCOOP-like motifs essential to trigger the MIK2-dependent immunity. However MIK2 has also been suggested to be the receptor of the Arabidopsis PROSCOOP family members including SCOOP4, 6, 8, 10, 12, 13, 14, 15 20 and 23^19,20^. In this study we report a novel role for MIK2 and SCOOP10/12 peptides in regulating leaf senescence. Our findings show that MIK2 has a negative regulatory role in both age-dependent and dark-induced leaf senescence. Mutants with disrupted MIK2 function exhibited precocious senescence, whereas transgenic plants expressing MIK2 under a senescence-induced promoter showed delayed senescence phenotypes. We also found that plants lacking PROSCOOP10 displayed a delayed senescence, while exogenous application of synthetic SCOOP10 peptides in detached leaves accelerated senescence in a MIK2-dependent manner. By contrast, loss-of-function *scoop12* mutants exhibited accelerated leaf senescence phenotypes and SCOOP12 peptides inhibited senescence in a MIK2-dependent manner. Both SCOOP10 and SCOOP12 peptides undergo a competitive binding regulation with the extracellular domain of MIK2, suggesting that MIK2 functions as a receptor for both senescence-promoting (SCOOP10) and senescence-inhibiting (SCOOP12) peptides. Our results suggest the presence of two modules (SCOOP10-MIK2 and SCOOP12-MIK2) operating antagonistically to fine-tune senescence in Arabidopsis leaves.

## Results

### MIK2 negatively regulates leaf senescence

To identify new receptor-like kinases (RLKs) regulating leaf senescence, we conducted a phenotypic analysis using a collection of over 200 T-DNA mutants obtained from the Arabidopsis Biological Resource Center (ABRC). These mutants had mutations in RLK genes (Table S1). Here, we report that plants lacking *MIK2* showed accelerated dark and age-induced leaf senescence. Two independent T-DNA insertion lines, *mik2-1* (SALK_061769) and *mik2-2* (CS419958) (Fig. S1A) were used for loss-of-function analysis. When compared to the corresponding wild type plants (Col-0, hereon WT), detached leaves from the *mik2* mutants exhibited accelerated senescence after 6 days of darkness (Fig. 1A, B). Lack of *MIK2* also caused premature leaf senescence in plants grown under controlled laboratory conditions (Fig. 1C, D). Consistently, rosette leaves of *mik2* plants exhibited lower chlorophyll (Chl) content, lower photochemical efficiency of photosystem II (PSII) (Fv/Fm ratio) and higher membrane ion leakage than that of WT (Fig. 1E). We found that *MIK2* is mainly expressed in aging leaves (Fig. S1B, C) and MIK2 proteins were confirmed to be localized on the plasma membrane (Fig S1D). The premature senescence phenotype observed in the *mik2* mutants was reversed to the WT phenotype by introducing the *MIK2* coding region under the control of a 2200 bp *MIK2* native promoter (Fig. 1F, G). Two individual transgenic lines with senescence induced-expression of *MIK2* driven by *pSAG12* (Senescence-specific promoter)^22,23^ were generated and displayed delayed leaf senescence with increased chlorophyll contents at the fourth, fifth and sixth rosette leaves in comparison to WT plants (Fig. 1H, I), highlighting that MIK2 presence was sufficient to inhibit leaf senescence supporting the role of MIK2 in the regulation of leaf senescence.

**Figure 1.**
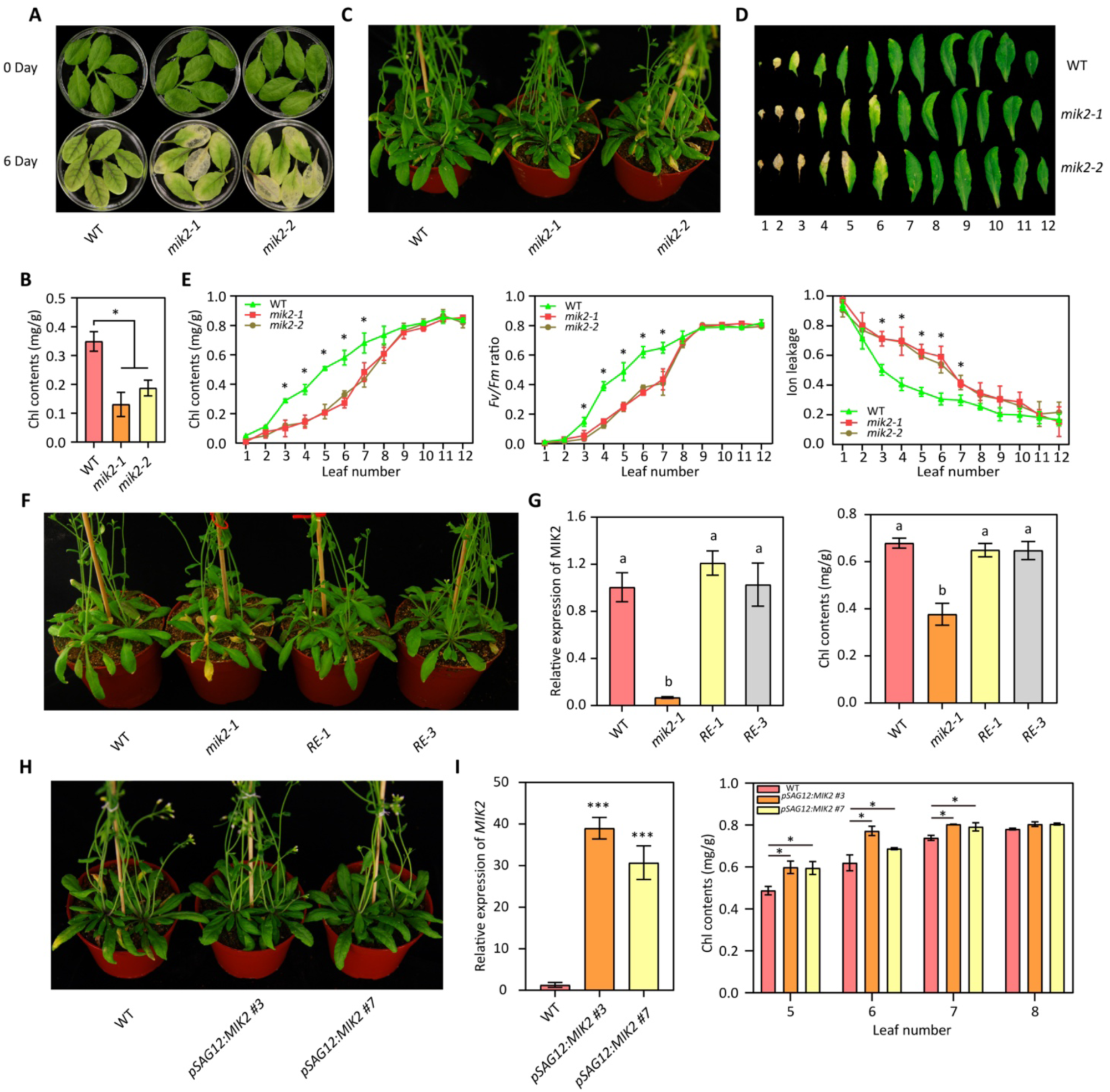
MIK2 negatively regulates both age-dependent and dark-induced leaf senescence. (A) Leaf senescence phenotype was assessed in detached 6^th^ rosette leaves obtained from 4-week-old *mik2-1* and *mik2-2* mutants and Col-0 (WT) plants after 6 days of dark treatment. Representative results from 3 independent experiments are presented. (B) Chlorophyll (Chl) contents were analyzed in the 6^th^ detached leaves as indicated in (A). Three biological replicates were included in each experiment and each replicate contained 6 individual leaves. (C) The adult plant phenotype of *mik2* mutants and WT plants was observed after 40 days of growth under long-day conditions. (D) Senescence progression was evaluated in the first twelve leaves isolated from the plants depicted in (C). (E) Measurements of Chl contents, PSII efficiency expressed as Fv/Fm ratio, and ion leakage were performed in adult plants of WT, *mik2-1*, and *mik2-2* mutants. Three biological replicates (each with 3 leaves) were analyzed for each experiment. (F) Leaf senescence phenotype was examined in two independent homozygous transgenic lines expressing *pMIK2*:*MIK2* (RE-1 and RE-3) in *mik2-1* mutant background. (G) Analysis of *MIK2* expression and measurement of Chl contents were conducted in the 6^th^ rosette leaf of the *pMIK2*:*MIK2 mik2-1* transgenic plants. Three biological replicates were used for all the physiological measurements. (H) Phenotypical analysis conducted on *pSAG12*:*MIK2*-expressing plants grown for 45 days. Two independent lines, *pSAG12*:*MIK2* #3 and *pSAG12*:*MIK2* #7, were tested. Three biological replicates were included in each experiment. (I) Relative expression of *MIK2* and Chl contents were analyzed using the 5^th^, 6^th^, 7^th^, and 8^th^ rosette leaves of the plants described in (H). Error bars represent the means ± standard deviations (SD) of three biological replicates for each experiment. Asterisks indicate statistically significant differences compared to WT based on ANOVA analysis (*P ≤ 0.05, ***P ≤ 0.01). Different letters (G) indicate statistically significant differences compared to WT as determined by one-way ANOVA analysis (P ≤ 0.05).

### SCOOP10 positively regulates leaf senescence

Because MIK2 was recently reported to function as receptor of the phytocytokine SCOOPs^19,20^ we tested whether SCOOP family peptides played a regulatory role in leaf senescence. First *PROSCOOP* expression was analyzed during different leaf developmental stages (Fig. S2A). Several *PROSCOOPs* showed a significant change in expression during leaf senescence and the exogenous application of several synthesized SCOOP peptides caused changes in senescence progression in detached leaves (Fig. S2B). Among those, SCOOP10 peptide (whose expression was found upregulated in the early stages of senescence process) treatments led to a premature senescence phenotype (Fig. 2A, B). Similarly, exogenous application of synthesized SCOOP10 peptides on intact rosette leaves caused significantly accelerated leaf senescence which was associated with reduced chlorophyll content and Fv/Fm ratio (Fig. S3A, 2B). To better understand the role of SCOOP10 in leaf senescence, two loss-of-function *proscoop10* T-DNA mutant lines (*proscoop10-1*, SALK_080439; and *proscoop10-2*, SALK_027949) were analyzed. Both lines displayed a delayed leaf senescence phenotype (Fig. 2C, D) with higher Chl content, Fv/Fm ratio and lower ion leakage (Fig. 2G, 2H, 2I). Transgenic *pSAG12: PROSCOOP10* lines in Col-0 (WT) showed an earlier senescence phenotype compared to the non-transformed WT plants (Fig. 2C-E). *pSAG12: PROSCOOP10* plants were in fact characterized by increased *PROSCOOP10* expression (Fig. 2E), yellower leaves (Fig. 2F), decreased Chl content, Fv/Fm ratio and increased ion leakage rate (Fig. 2G, 2H, 2I). Interestingly, *PROSCOOP10* transcripts were found significantly up-regulated in dark-treated leaves (Fig. S3C). Furthermore, detached leaves from *proscoop10* mutants displayed a delayed senescence phenotype after 8 days of dark treatments (Fig. S3D, E). To investigate whether MIK2 is required for PROSCOOP10-triggered leaf senescence, detached leaves of *mik2* mutants were treated with SCOOP10 synthetic peptides. Four days after treatments, premature leaf senescence was observed in the WT but not in *mik2* mutant alleles (Fig. 3A-C). *mik2* mutant lines were nonresponsive to SCOOP10 peptide application strongly suggesting that MIK2 is required for the perception of SCOOP10 and for the SCOOP10-mediated induction of leaf senescence. To confirm this, we generated a *proscoop10-1 mik2-1* double mutant line. The *proscoop10-1 mik2-1* plants were characterized by premature senescence phenotypes when compared to *proscoop10-1* or WT plants, likely reflecting the *mik2-1* mutant phenotype (Fig. 3D, E). Consistent with the premature senescence phenotype, *proscoop10-1 mik2-1* double mutant displayed lower Chl content and *SAG12* up-regulation when compared to that of the controls (Fig. 3F). Furthermore, a microscale thermophoresis (MST) assay was performed to determine the ability of MIK2 ectodomain (expressed in *Escherichia coli* cells) in binding the FAM labeled-SCOOP10 (5-FAM-SCOOP10) peptide (Fig. 3G). The biological functions of the 5-FAM-SCOOP10 peptides i.e., induce leaf senescence and/or root growth inhibition^19,20^ were confirmed to be similar to those triggered by the unlabeled peptides (Supplementary Fig. 4A, B). The results of the MST analysis indicated that the FAM-SCOOP10 peptide was able to bind to the ectodomain of MIK2 with a dissociation constant (Kd) of 13.50 ± 1.47 nM (Fig. 3G), which suggested a direct interaction between SCOOP10 and MIK2 *in vitro*. Taken together, the results show that SCOOP10 interact with MIK2 and that PROSCOOP10 mediates leaf senescence in a MIK2-dependent manner. These findings collectively indicate that SCOOP10 plays a positive regulatory role in both age-dependent and dark-induced leaf senescence.

**Figure 2.**
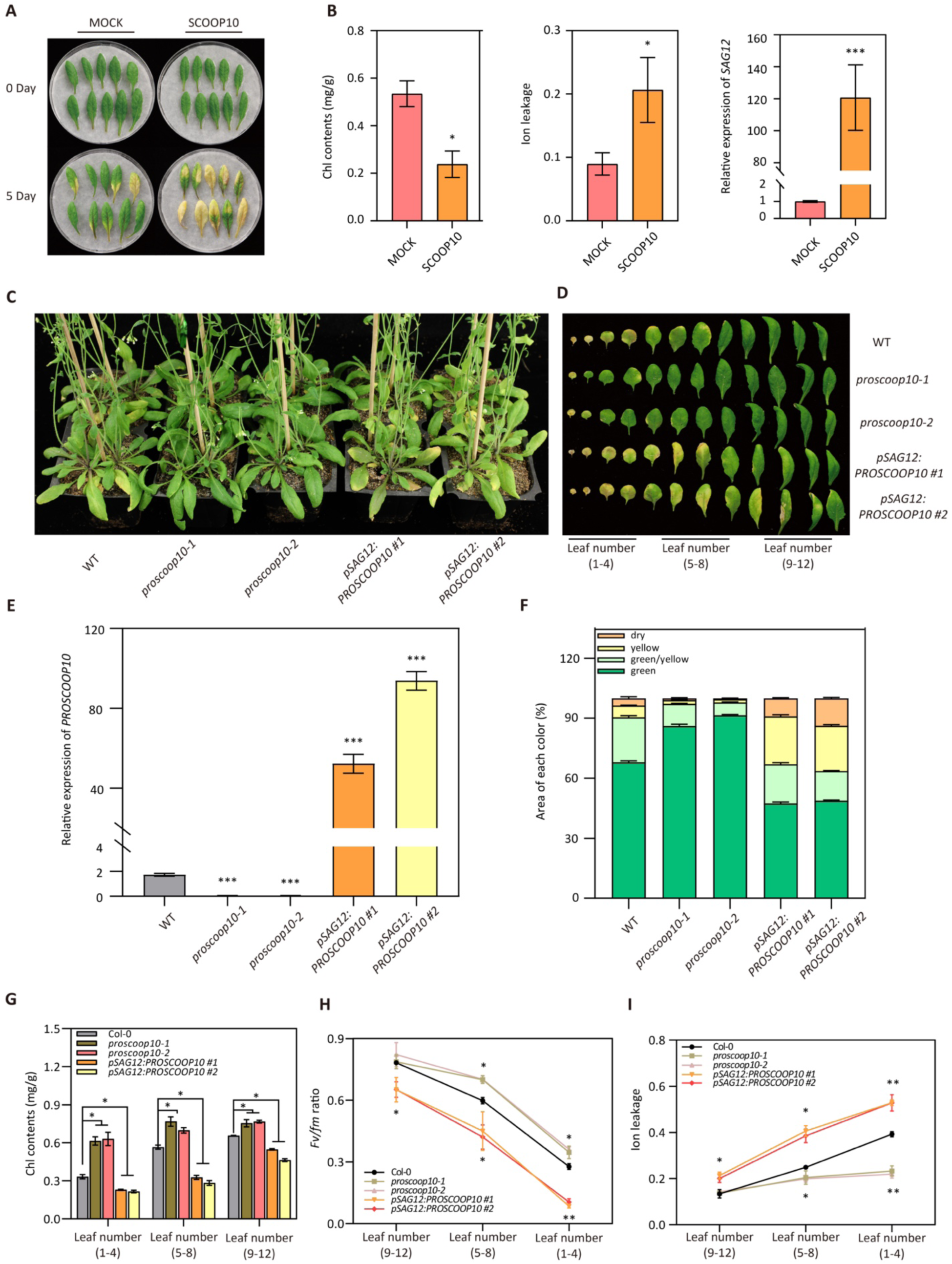
SCOOP10 positively regulates leaf senescence. (A) Leaf senescence was analyzed in the 5^th^ and 6^th^ rosette leaves derived from 4-week-old WT plants treated with chemically synthesized SCOOP10 peptides (1 µM) for 5 days. Representative results are shown from three independent experiments. (B) Chl contents, ion leakage, and *SAG12* expression were analyzed in the detached leaves shown in (A). (C and D) Rosette leaves of *proscoop10* mutants, *pSAG12*:*PROSCOOP10* #1 and #2, and corresponding WT were isolated and grouped into leaf numbers 1-4, 5-8, and 9-12. (E) *PROSCOOP10* expression in *proscoop10* mutants, *pSAG12*:*PROSCOOP10* #1 and #2, and corresponding WT. (F) Area of each color of *proscoop10* mutants and the *pSAG12*:*PROSCOOP10* #1 and #2 transgenic lines grown under long-day conditions for 45 days. (G) Chl contents, Fv/Fm ratio (H), and ion leakage (I) were measured in the detached leaves presented in (C and D). Error bars represent means ± SD of three biological replicates for each experiment. Ten, 3, and 4 leaves were used in measurements presented in B, F, and G-I, respectively. Asterisks indicate significant differences compared to WT, as determined by a two-way ANOVA analysis (*P ≤ 0.05, ***P ≤ 0.01).

**Figure 3.**
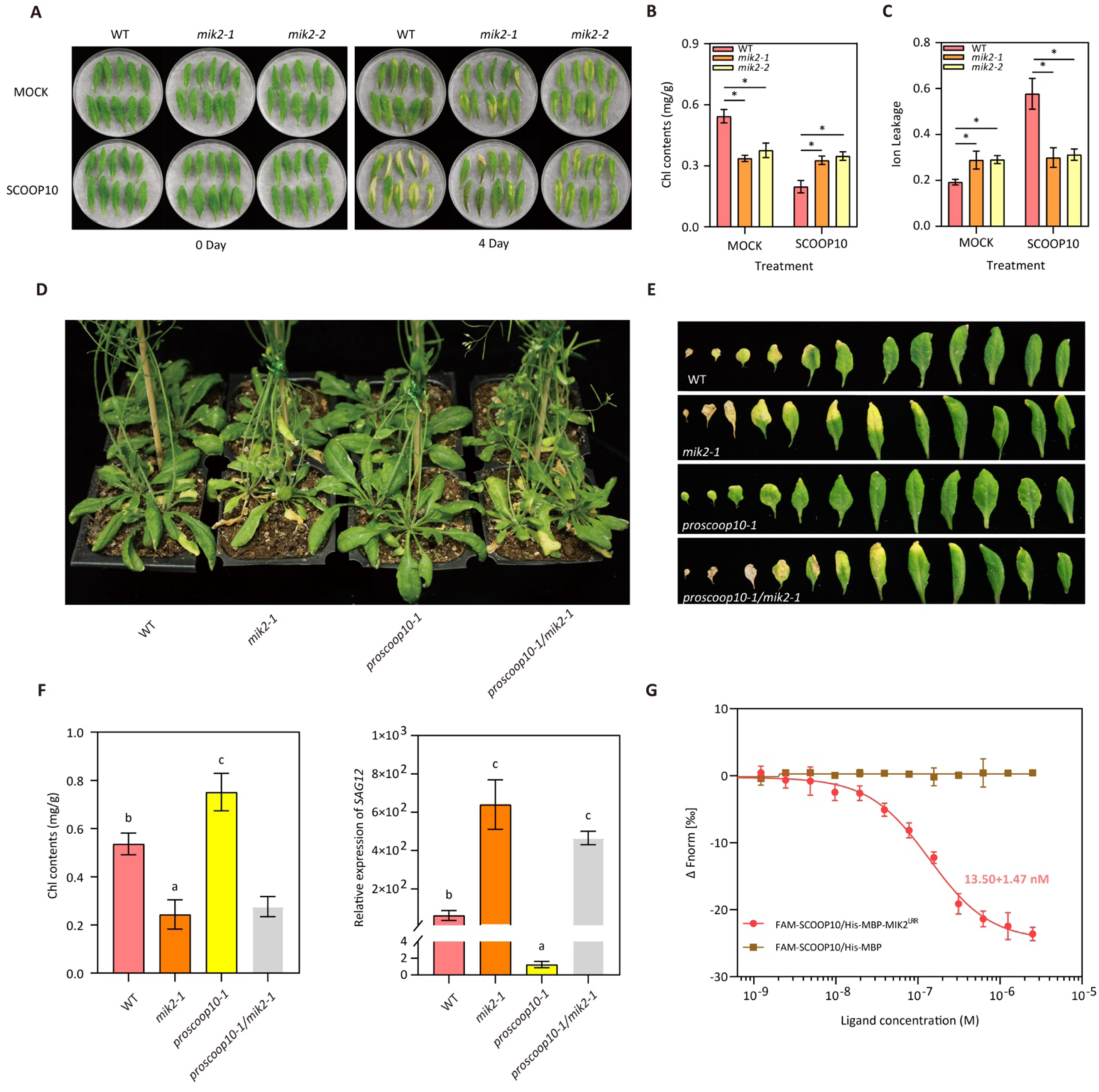
SCOOP10 regulates leaf senescence in a MIK2-dependent manner. (A) Leaf senescence was analyzed in the 5^th^ and 6^th^ rosette leaves derived from 4-week-old *mik2* mutants and corresponding WT plants upon treatments with synthesized SCOOP10 peptides (1 μM) for 2 and 4 days under light conditions. (B) Changes in Chl contents and (C) ion leakage of detached leaves shown in (A) after 4 days of treatments. Three biological replicates were included in each experiment and each replicate contained ten leaves. (D) Leaf senescence phenotype of 45-day-old *proscoop10-1 mik2-1* double mutants, *proscoop10-1* and *mik2-1* single mutants, and corresponding WT plants grown under long-day conditions. (E) Chl contents and transcript levels of *SAG12* measured in the 6^th^ rosette leaves isolated from the plants presented in (D). Six leaves were analyzed for each replicate. (F) Senescence phenotype of the first twelve rosette leaves of the plants in (D). (G) Determination of the binding affinity between SCOOP10 peptides with a fluorescent FAM label on the N-terminus and the ectodomain of MIK2 using microscale thermophoresis (MST). FAM-SCOOP10/MBP was used as a negative control in the binding assay. All results are presented as the mean ± SD of three independent experiments. Asterisks were used to indicate significant differences. Letters represent statistically significant differences compared to WT, as examined based on one-way ANOVA analysis (*P ≤ 0.05, ***P ≤ 0.01).

### SCOOP12 negatively regulates leaf senescence in a MIK2-dependent manner

Because SCOOP12 peptide was recently reported to directly bind the ectodomain of MIK2 *in vitro*^20^, we also studied the function of SCOOP12 in regulating leaf senescence. While the expression of *PROSCOOP12* was dramatically induced during aging (Fig. S2A), exogenous application of SCOOP12 peptides significantly inhibited senescence (Fig. 4A, B, C, D), suggesting that SCOOP12 might function as a negative regulator of senescence. Further analyses showed that detached leaves of *mik2* mutants were nonresponsive to the exogenous application of SCOOP12 peptide (Fig. 4E, F, G), indicating that SCOOP12-mediated inhibition of leaf senescence requires a functional MIK2. Furthermore, loss-of-function *proscoop12* mutant lines^24^ displayed premature senescence while senescence driven expression of *PROSCOOP12* (*pSAG12:PROSCOOP12* transgenic plants) led to delay in leaf senescence (Fig. 4H and I). To further investigate the genetic interaction between PROSCOOP12 and MIK2, we found that double mutants (*pSAG12:PROSCOOP12 #6 mik2-1* double mutants) displayed an earlier senescence phenotype comparable to that of the *mik2* loss-of-function mutants (Fig. 4H and I). Consistent with the previous findings^20^, the MST results indicated a direct binding between ectodomain of MIK2 and SCOOP12 with a Kd of 21.98 ± 3.02 nM (Fig. 4J). Together, these data suggest that the senescence inhibiting effect of SCOOP12 is genetically dependent on MIK2. These findings strongly support the hypothesis that not only SCOOP10 but also SCOOP12 interacts with MIK2 to modulate senescence progression.

**Figure 4.**
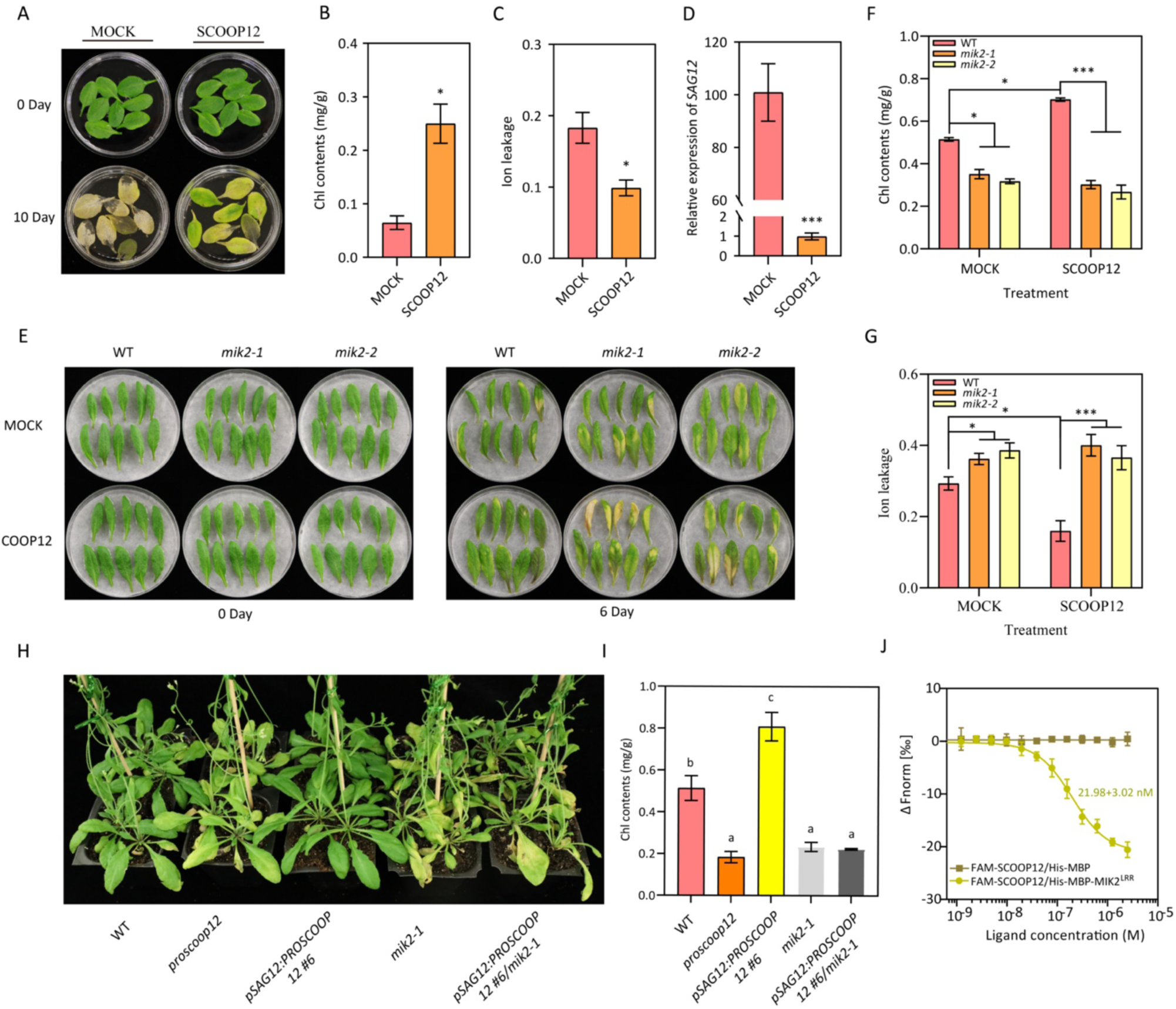
SCOOP12 negatively regulates leaf senescence in an MIK2-dependent manner. (A) Leaf senescence phenotype was analyzed in the 5^th^ rosette leaves from 4-week-old plants upon treatments with synthesized SCOOP12 peptides (1 μM) for ten days. (B-D) Chl contents (B), ion leakage (C) and *SAG12* expression (D) were measured in detached leaves shown in (A) ten days after treatments. Three biological replications were performed with 8 leaves per replicate. (E) Senescence phenotypes analyzed in the 5^th^ and 6^th^ rosette leaves from 4-week-old *mik2* mutants and corresponding WT treated with synthesized SCOOP12 peptides (1 µM) for 6 days under light condition. Chl contents (F) and ion leakage (G) were measured in detached leaves shown in (E) Ten leaves were measured in each replicate. (H) Senescence phenotype was analyzed in 6-week-old *pSAG12*:*SCOOP12* #6 in the *mik2-1* background, *pSAG12*:*SCOOP12* #6, *proscoop12*, *mik2-1* and WT grown under long-day conditions. (I) Chl contents of the 6^th^ rosette leaves collected from plants shown in (H). Three biological replications were included in each experiment. (J) MST analysis of binding affinity between SCOOP12 peptides and the ectodomain of MIK2. Error bars represent SD of three independent experiments. All results are presented as the mean ± SD of three independent experiments. Asterisks was used to indicate significant differences. Different letters represent statically significant differences to WT based on a one-way ANOVA analysis (*p < 0.05 and ***p < 0.01).

### SCOOP10 and SCOOP12 act antagonistically in regulating leaf senescence through MIK2

To investigate the interactions between SCOOP10 and SCOOP12 in the regulation of the MIK2-dependent leaf senescence, WT detached leaves were treated with SCOOP10 and SCOOP12 peptides simultaneously. Interestingly, SCOOP10-triggered leaf senescence was alleviated by SCOOP12 peptide application (Fig. 5A, B, C), suggesting an antagonistic effect between the two peptides. To explore this antagonistic mechanism further, a sequential treatment assay was designed, where detached leaves were initially pretreated with one peptide for a period of 2 days, followed by the application of the other peptide for a subsequent 4-day treatment (Fig. 5D). As a result, we observed that leaves treated with SCOOP12 for 2 days prior to the application of SCOOP10 exhibited a senescence phenotype similar to the mock treatment, in contrast to the significant senescence phenotypes displayed in leaves treated with SCOOP10 alone (Fig. 5E, F). On the other hand, when detached leaves were pretreated with SCOOP10 peptides for 2 days before the application of SCOOP12, they exhibited a delayed senescence phenotype similar to the leaves treated with SCOOP12 alone (Fig. 5E, F). The results suggest a dominant effect of SCOOP12 over SCOOP10 in the regulation of senescence. Because SCOOP12 and SCOOP10 are both recognized by MIK2, the SCOOP peptides might compete for interaction with the MIK2 receptor and SCOOP12 might have advantage in this competition. To test this hypothesis, an MST assay was set up to analyze the MIK2 binding competition between SCOOP10 and SCOOP12 and the results indicated that SCOOP12 peptides inhibited the interaction between SCOOP10 peptides and MIK2 with an inhibition constant (Ki) of 40.15 nM (Fig. 5G).

**Figure 5.**
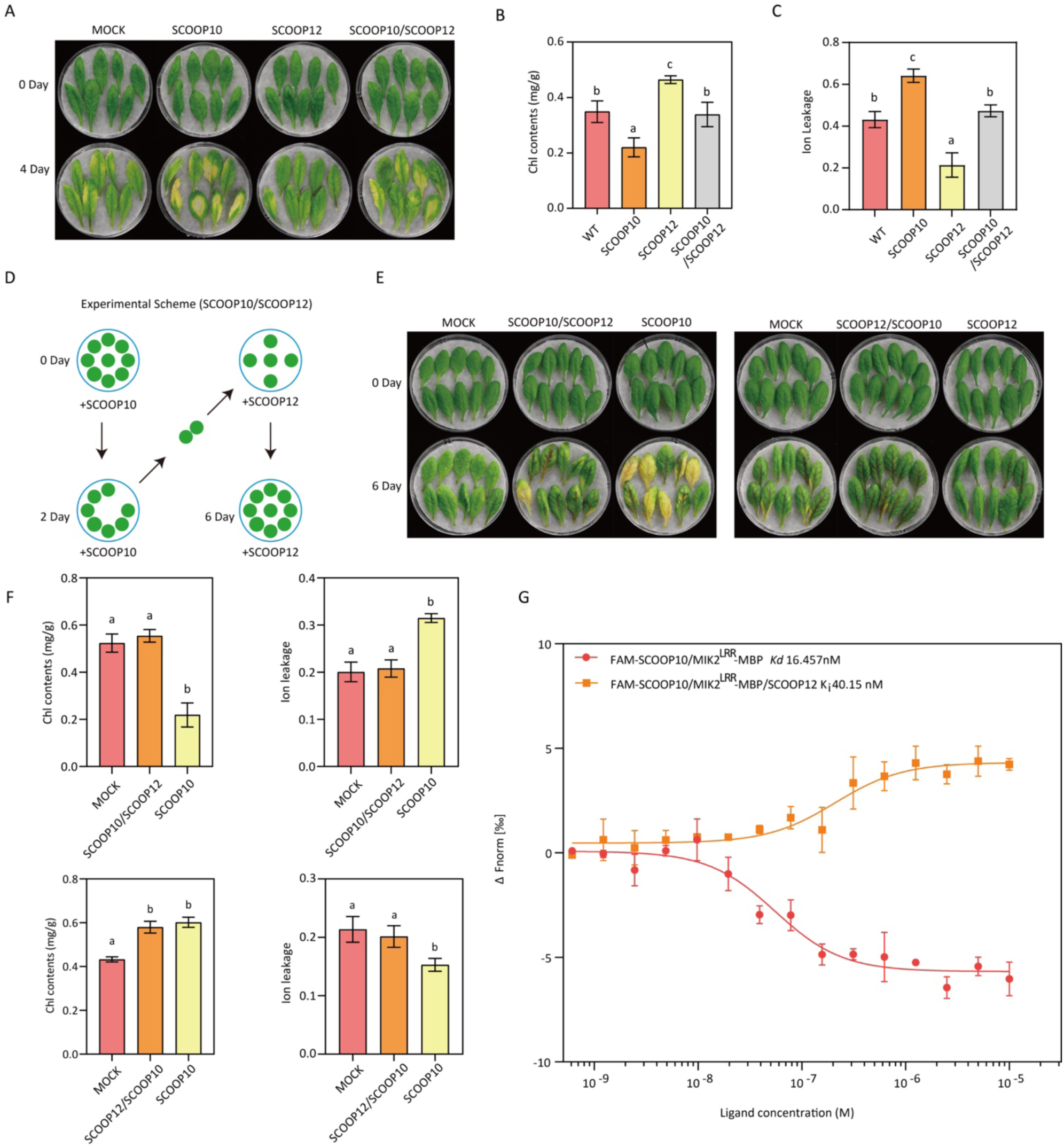
SCOOP12 competes with SCOOP10 for MIK2 interaction in regulating leaf senescence. (A) Senescence phenotype of the 6^th^ rosette leaves from 30-day-old WT plants were analyzed upon treatments with 1 µM SCOOP10, SCOOP12, SCOOP10 plus SCOOP12 (SCOOP10/SCOOP12) or mock (ddH2O) after 4 days under long-day conditions. Chl contents (B) and ion leakage (C) were measured in detached leaves as shown in (B). Three biological replicates were performed with each containing 8 leaves. (D) The experimental scheme for competitive assay between SCOOP12 and SCOOP10 peptides in regulating leaf senescence. The 5^th^ and 6^th^ rosette leaves from 4-week-old plants were first pretreated by SCOOP12 peptides for 2 days, then transferred to ddH2O containing SCOOP10 peptides and incubated for 4 additional days under long-day conditions. (E) Senescence phenotype analyzed in detached leaves pretreated either with SCOOP10 or SCOOP12 peptides for 2 days and then treated in SCOOP12 or SCOOP10 peptides for 4 additional days. (F) Chl contents and ion leakage measured in detached leaves shown in (E). Three replicates were performed with each containing at least 8 leaves. (G) Competitive inhibition analysis of SCOOP12 and FAM-SCOOP10 in binding the MIK2 ectodomain was performed with MST. Bars indicate means ± SD calculated by three biological replicates. Different letters represent statically significant between treatments and controls based on a one-way ANOVA analysis (P < 0.05).

*PROSCOOP10* expression is upregulated during the early senescence stage while *PROSCOOP12* is only induced during the late senescence stage (Fig. 6A). In addiction the application of exogenous SCOOP10 peptides was found to induce the expression of *PROSCOOP12* within a time frame of 10 hours (Fig. 6B). This observation suggests the presence of a negative feedback mechanism, where SCOOP10 and SCOOP12 regulate each other as potential modulators of the signaling pathway fine-tuning the senescence progression. The sequential and antagonistic interaction between SCOOP10-MIK2 and SCOOP12-MIK2 suggests the presence of a finely balanced regulatory mechanism (schematized in Fig. 6C) creating a knot-like structure that ensures a timely progression of leaf senescence.

**Figure 6.**
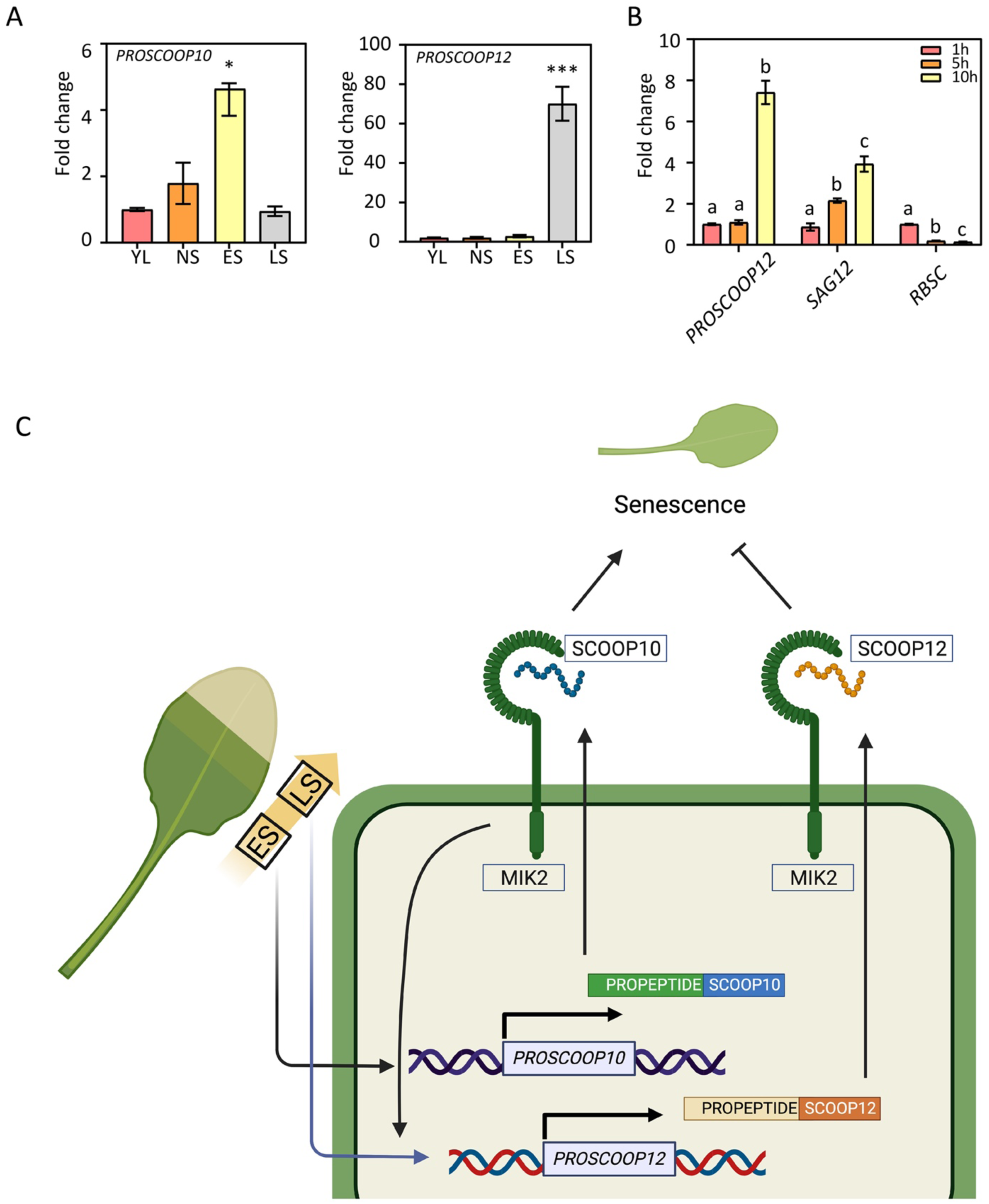
A model of the antagonistic functions of SCOOP10 and SCOOP12 in regulating leaf senescence by competitively binding to MIK2. (A) Fold changes in the expression of *PROSCOOP10* and *PROSCOOP12* at different stages of leaf senescence. Three independent experiments were performed. The bars indicate means ± SD calculated by three biological replicates, and asterisks represent statically significant differences to YL, tested by a one-way ANOVA analysis (*p < 0.05 and ***p < 0.01). Young leaf (YL), fully extended non-senescent leaf (NS), early senescent leaf (ES) with approximately 25% yellowing area, and late senescent leaf (LS) with more than 50% yellowing area. (B) Expression analysis of *PROSCOOP12*, *SAG12* and *RBCS* after treatments with synthetic SCOOP10 (1 µM) peptides for 1, 5 and 10 hours. Three biological replicates were analyzed in each experiment, and different letters represent statically significant differences to control (1 h), according to Student’s T-test P < 0.05. (C) Hypothetical model based on our findings shows the possible mechanism through which SCOOP10 and SCOOP12 regulate leaf senescence by competitive interaction with MIK2. The transcriptional activation of *PROSCOOP10* and *PROSCOOP12* is regulated by age. During the early stage of leaf senescence *PROSCOOP10* accumulates and SCOOP10 binding to MIK2 triggers senescence, likely functioning as a positive regulator of leaf senescence. SCOOP10 recognition boosts the expression of *PROSCOOP12* likely resulting to an increased SCOOP12 synthesis and consequent inhibition of leaf senescence. (ES) Early senescent leaf and (LS) Late senescent leaf.

## Discussion

As a genetically regulated age-dependent process, leaf senescence involves signaling events that are triggered by various endogenous and environmental factors. The integration of these factors determines the progression of the so-called “senescence syndrome”^25,26^. In this study, we discovered a novel senescence-regulating signaling module involving the receptor-like kinase MIK2 and two of the SCOOP peptides. MIK2 is a member of the subfamily XI of Leucine-Rich Repeat (LRR) Receptor like Kinases (RLK) and was firstly identified to control pollen tube guidance and plant fertilization by cooperating with MIK1 and MALE DISCOVERER1 (MDIS1) for recognition of the peptide LURE1^27,28^. Later, through a GWAS screening MIK2 was associated to salt stress responses. Arabidopsis ecotypes displaying an increased expression of *MIK2* (or *LRR-KISS*) were characterized by enhanced tolerance to salt stress, while knock-down line HR-5 with a reduced expression of *MIK2* displayed increased salt susceptibility^29^. MIK2 was also linked to cell wall integrity^30,31^and disease resistance^30^. MIK2 was reported to function in elicitor perception in response to infection by Fusarium spp. fungi^30,31^. It was found that exogenous peptides corresponding to Fusarium-derived SCOOP-like motifs could trigger MIK2-dependent immunity against Fusarium. MIK2 was thus suggested as a receptor of SCOOP family members in Arabidopsis and was shown to interact directly with SCOOP12. Loss-of-function in *proscoop12* or *mik2* impaired plant resistance against pathogens and *mik2* was insensitive to the SCOOP12-dependent signaling activation^20^. In this study we uncovered the regulatory role of MIK2 during leaf senescence. Similar to its involvement in stress responses, increased expression of *MIK2* leads to a delay in senescence while *mik2* mutant plants exhibit premature leaf senescence (Fig. 1). Because MIK2 have been previously implicated in responses to abiotic stresses^29^, pathogen infection^30^, it is plausible that MIK2 serves as a critical hub in mediating signaling pathways triggered by diverse physiological factors, likely affecting plant senescence. Similarly small peptides have been characterized to function in various developmental and stress response processes, usually through receptor-like kinases^32^. In most cases binding of the peptide ligands affect the downstream signaling in a positive or negative manner^33^. Here we report that SCOOP10 and SCOOP12 peptide treatments caused the opposite phenotype of leaf senescence both in a MIK2-dependent manner (Fig. 3, 4). It is possible that in the SCOOP10/SCOOP12-MIK2 regulatory pathway described in this study, PROSCOOPs are expressed during different stages of senescence likely controlling senescence, in temporal manner. In fact, we found that *PROSCOOP10* is highly expressed during the early stages of senescence (Fig. 6, Fig S2), likely promoting the initiation of senescence. But as senescence progresses, SCOOP10 signaling triggers the expression of *PROSCOOP12* likely altering its progression (Figure 6B). This sequential activation of SCOOP12 suggests a dynamic interplay between SCOOP10 and SCOOP12, mediated by MIK2, as an intricate regulatory mechanism during leaf senescence. Binding of SCOOP10 peptides to MIK2 can be displaced by SCOOP12 (Fig 5G), leading to a decrease in *PROSCOOP10* expression. As senescence progresses, SCOOP12 becomes the dominant peptide that binds to MIK2. Peptide ligands from the same family have been reported to be able to bind to the same receptor leading to opposite responses ^19,34^. Similar antagonistic regulations have been reported recently for the RAPID ALKALINIZATION FACTOR (RALF) peptide family. During pollination RALF4/19 and RALF34 antagonistically regulate pollen tip integrity to avoid premature burst^35^. RALF23 and RALF33 peptides compete with the POLLEN COAT PROTEIN B-class peptides (PCP-Bs) in binding to the *Catharanthus roseus* RLKs ANJEA–FERONIA (ANJ–FER) receptor complex, in regulating reactive oxygen species (ROS) production during the stigma and pollen hydration^36^. Here, we envision a mechanism where the activation of the MIK2-mediated senescence pathway by SCOOP12 might acts as a safeguard, preventing senescence from progressing too rapidly and ensuring the orderly progression of nutrient recycling (Figure 6). The SCOOP-MIK2 module represents a novel mechanism in plant senescence, where antagonistic peptide competition for the same receptor to fine-tune the control of leaf senescence. Our discovery highlights the intricate and dynamic nature of peptide signaling in plant development and provides insights into the regulatory networks governing leaf senescence.

## Materials and methods

### Plant growth and detached leaf treatments

*Arabidopsis thaliana* ecotype Columbia-0 and mutants used in this study include: *mik2-1* (SALK_06269), *mik2-2* (CS419958), *proscoop10-1* (SALK_080439), *proscoop10-2* (SALK_027949) that were obtained from the Arabidopsis Biological Resource Center (ABRC). The *proscoop12* CRISPR-Cas9-generated mutants were kindly provided by Dr. Elia Stahi at University of Lausanne, Switzerland^24^. Imbibed seeds were incubated at 4°C in a refrigerator for 2 days before sowing on the soil mix (3 parts commercial soil: 1 parts vermiculite) and grown under the long-day-condition (16/8 h) in a growth chamber with 150 μmol m^-2^ sec^-1^ light intensity. For the senescence phenotyping of detached leaves, the fifth and sixth rosette leaves of 25-day-old plants were removed and placed in Petri dishes with the adaxial side facing up. The Petri dishes were sealed with parafilm to prevent water loss, and the leaves were incubated with deionized water. The application of exogenous peptides followed the previously established method^12,13^. For dark-induced leaf senescence assays, detached rosette leaves were incubated in deionized water placed in the dark condition for 6 days. At least three biological replicates were included for each assay. For senescence phenotype analysis under normal growth conditions, homozygous plants were grown alongside with Col-0. All data including plant phenotypes and gene expression presented in this study were analyzed with Graphpad Prism 9 and at least there biological replicates were included in each experiment. Chlorophyll content, photochemical efficiency of PSII (Fv/Fm), and ion leakage were determined as previously described ^12,14,37^

### Generation of transgenic plants

The 2200 bp *MIK2* promoter (*pMIK2*) and the 2000 bp *SAG12* promoter (*pSAG12*) were amplified from Col-0 genomic DNA. The Coding Sequences (CDS) of *MIK2*, *PROSCOOP10* and *PROSCOOP12* were cloned via RT-PCR. The cloned fragments were inserted into the pEASY-Blunt vector (TransGen Biotech) to generate sub-clone vectors *pMIK2*-Blunt, *pSAG2*-Blunt, *gMIK2*-Blunt, *gSCOOP10*-Blunt, and *gSCOOP12*-Blunt, which were confirmed by sequencing. For complementation test, *pMIK2* and *MIK2* CDS from *pMIK2* and *gMIK2*-Blunt were inserted into the *pPZP211* vector separately at the *EcoRI* and *SacI* sites to generate the *pMIK2:MIK2* construct by Infusion (Vazyme Biotech Co., Ltd). For *SAG12*-induced expression, *pSAG12* and the CDS of *MIK2*, *PROSCOOP10* as well as *PROSCOOP12* were cloned into *pPZP211* to obtain *pSAG12:MIK2*, *pSAG12:PROSCOOP10* and *pSAG12:PROSCOOP12*. For subcellular localization assays, the *MIK2* CDS without stop codon fused with GFP was cloned into pCHF3 to generate pCHF3::MIK2-GFP vector by Infusion upon digestion at the *Sac I* site. For generation of pMIK2::MIK2-GFP vector, the sequence of *pMIK2* and *MIK2-GFP* were amplified using *pMIK2*-Blunt and pCHF3::MIK2-GFP vectors as template, and then inserted into the pZP211 backbone. All primers used in this study were synthesized by Personal Biotechnology Co.,Ltd (Shanghai, China) and the sequences are listed in Table S2. The recombinant vectors described above were transformed into *Agrobacterium tumefaciens* strain GV3101 for subsequent transformation of Arabidopsis plants via the floral dipping method. For the screening for transgenic plants, seeds were surface-sterilized with 75% ethanol for 5 min, washed with 100% ethanol for 1 min and sown on half-strength Murashige and Skoog medium (1/2MS) containing appropriate antibiotics, pH 5.8, followed by 2 days’ incubation in the dark at 4°C. Ten days after germination, plants were transplanted to the soil mix in the growth chamber and grown under the conditions described above. Genotyping analyses were performed with leaf tissues from plants grown in soil using gene-specific primers (Table S2).

### Gene expression analysis

Total RNA was isolated from Arabidopsis leaves with the RNAiso Plus (TAKARA), and reverse transcribed by the PrimeScript™ RT reagent Kit (TAKARA). qPCR reactions were implemented in an Applied Biosystems 7500 Real-Time PCR system with the SYBR® Premix Ex TaqTM II (TAKARA). *ACTIN* was used as the reference gene for qPCR analyses. Primers used for qPCR are listed in Table S2.

### Protein expression and purification

To express MIK2 protein in the soluble form, the truncated sequence of MIK2 (MIK2 ^LRR^) expressing only the ectodomain (residues 45–707) was amplified from the *gMIK2*-Blunt subclone vector and inserted into the *pET41* vector, fused with a 6 × His-MBP tag. Then, the ectodomain of MIK2 (MIK2^LRR^) in the absence of the putative N-terminal signal peptide was expressed in *Escherichia coli* strain BL21 (TransGen). Cell cultures were grown to 0.6 OD600 and then induced with 0.3 mM isopropyl-β -d-thiogalactopyranoside (IPTG) for 30 h at 13°C. The mixture was then centrifuged at 12 000 rpm to separate the cells from the medium. The pellets were resuspended with the binding buffer (20 mM Phosphate buffer, 500 mM NaCl, 50 mM Imidazole, pH 7.4). The cells were sonicated on ice for 30 min in the lysis buffer (0.2 mg/ml lysozyme, 20 μg/ml DNAse, 1 mM MgCl2, 1 mM PMSF). After centrifugation (12000 rpm for 30 min at 4 °C), the supernatant was collected for purification. His-MBP-tagged MIK2 ^LRR^ isolation was performed using the Mag-Beads His-Tag Protein purification Kit (Sangon Biotech) following the instructions of the manufacturer.

### Microscale thermophoresis (MST) analysis

A Monolith NT.115 system (Nanotemper Technologies) was used to perform MST assays. To measure the binding affinity with MIK2^LRR^, SCOOP10 and SCOOP12 peptides were labeled with fluorescent 5-FAM at the N-termini (GenScript, China) and the final working concentration of labelled peptides was adjusted to 0.05 μM with ddH2O. The concentration of purified MIK2^LRR^ was measured with the BCA Protein Quantification Kit (Vazyme) and then diluted with ddH2O to 20 μM. To analyze the binding affinity between 5-FAM-SCOOP10/12 and MIK2^LRR^, the purified MIK2^LRR^ was incubated with FAM-SCOOP10 or FAM-SCOOP12 for 10 min at room temperature and then loaded into silica capillaries (Monolith™ NT.115 Standard Treated Capillaries, MO-K002). MST assays were performed according to protocols provided by NanoTemper Technologies (20% LED power and 40% MST power). MST results were analyzed using the Nanotemper analysis software (MO. Affinity)^38^. The competitive binding analysis were performed as described previously^36^. To analyze the competitive binding between SCOOP12 and SCOOP10 to MIK2, 0.5 µM FAM-SCOP10 peptide and 15 µM purified MIK2^LRR^ were mixed and incubated for 10 min at room temperature in the darkness. A range of concentrations of SCOOP12 peptides were introduced to the FAM-SCOOP10/MIK2 LRR mixture, and the mixture was co-incubated for 10 minutes. The mixtures were subsequently loaded into silica capillaries to conduct MST assays using 20 % LED power and 20 % MST power. The dissociation constant (Ki) was determined using the Ki Finder Tool available on the NanoTemper Technologies website (http://www.nanotemper technologies.com/get-it-all/ tools/ki-finder/).

### Peptides synthesis

All peptides used in this study were synthesized by GenScript (China) with >98% HPLC purity. The peptide sequences are listed in Table 3 referring to paper published previously (Hou et al., 2021 and Rhodes et al., 2022).

### Confocal microscopic analysis

To observe protein subcellular localization, MIK2 fused with GFP driven by the 35S promoter was transiently expressed in *Nicotiana benthamiana*. Images were obtained at 488 nm laser excitation using a confocal microscope (TCS-SP8, Leica, Wetzlar, Germany). The images were processed with the Leica Application Suite X (LAS X) software.

## Author contributions

Y.G. and N.G.B. conceptualized and designed the study. Y.G. C.T., N.G.B. and W.L. supervised Z.Z who performed and collected all the data presented in this manuscript. Z.Z and N.G.B. drafted the manuscript, Y.G., C.T. edited the manuscript. All authors critically reviewed the manuscript and approved it for publication.

## Acknowledgements

We thank Dr. Elia Stahi at University of Lausanne and the Arabidopsis Biological Resource Center (ABRC) for kindly providing the *proscoop12* mutants. This work was supported by the Agricultural Science and Technology Innovation Program, Chinese Academy of Agricultural Sciences (ASTIP-TRI02 to Y.G.), the National Natural Science Foundation of China (32270332 to Y.G. and 31970204 to W.L.), the Graduate School of Chinese Academy of Agricultural Sciences (Z. Z), Wageningen University Joint PhD Programme and the European Research Council (ERC) under the EU Horizon 2020 Research and Innovation Programme (grant agreement 724321 to C.T.).

## Supplemental Figure Legends

**Figure S1.**
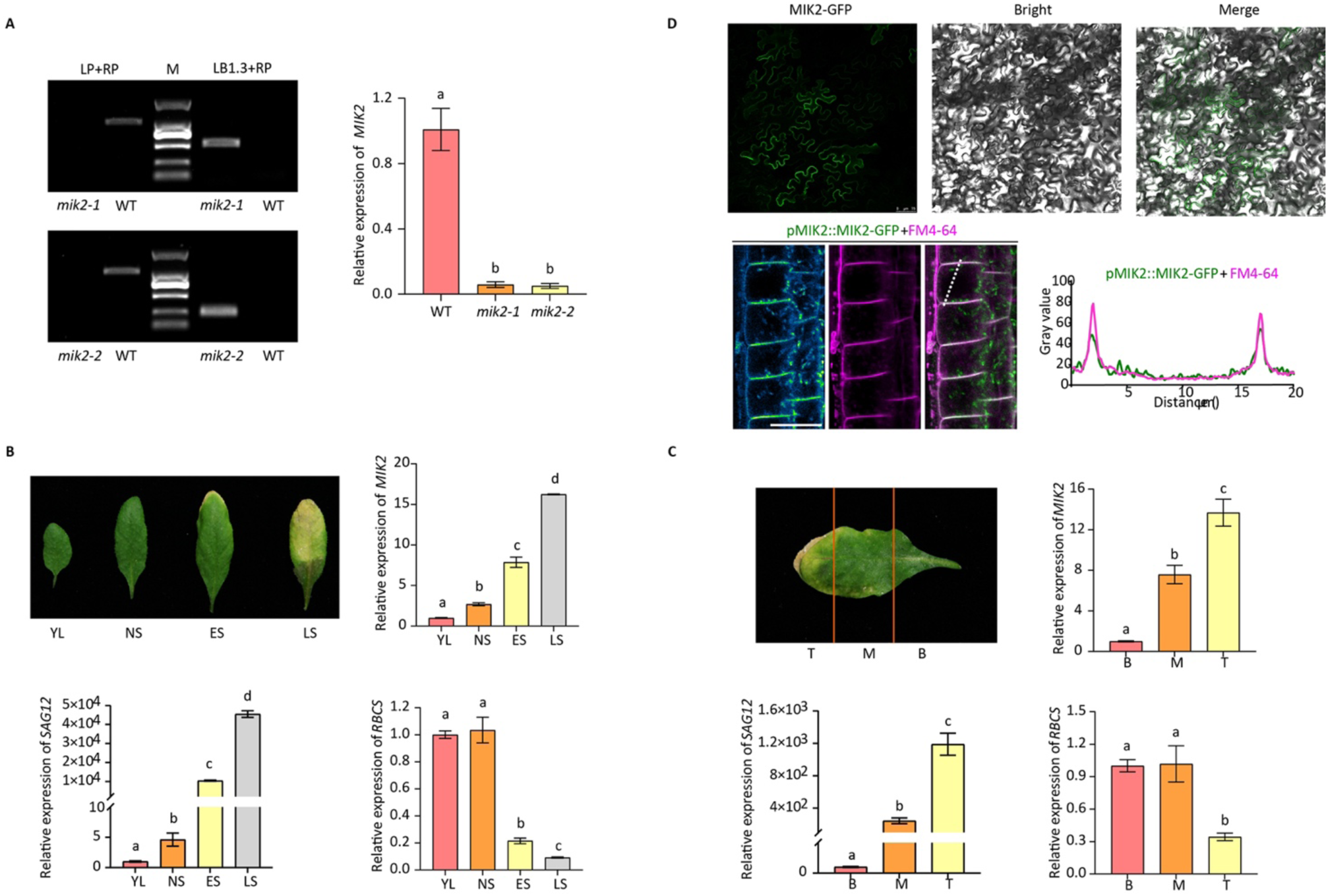
(A) Genotyping of T-DNA insertion mutants (*mik2-1* and *mik2-2*) and analysis of transcript levels using qRT-PCR. (B) Schematic representation of different senescence stages, along with the relative expression levels of *MIK2*, *SAG12*, and *RBCS* determined by qRT-PCR in the indicated leaves. The leaves are categorized as young leaf (YL), fully extended non-senescent leaf (NS), early senescent leaf (ES), and late senescent leaf (LS). The analysis was performed in three independent experiments. (C) Schematic representation of Arabidopsis 6^th^ rosette leaves from 5-week-old plants is presented, along with the expression analysis of *MIK2*, *SAG12*, and *RBCS* in different parts of a leaf, including the base (B), middle (M), and tip (T). The experiment was replicated three times. (D) Subcellular localization of the GFP-MIK2 fusion protein was detected in tobacco leaves (*35S:MIK2-GFP,* upper panel) and in Arabidopsis roots (*pMIK2*:*MIK2*-*GFP mik2-1*, lower panel) by using confocal microscopy. GFP-MIK2 co-localizes with the PM dye FM4-64 in *pMIK2*:*MIK2*-*GFP mik2-1* roots. The scale bar represents 75 µm. Bars represent means ± SD of three biological replicates. Letters represent statistically significant difference to control (YL) or WT based on a one-way ANOVA analysis (P < 0.05).

**Figure S2.**
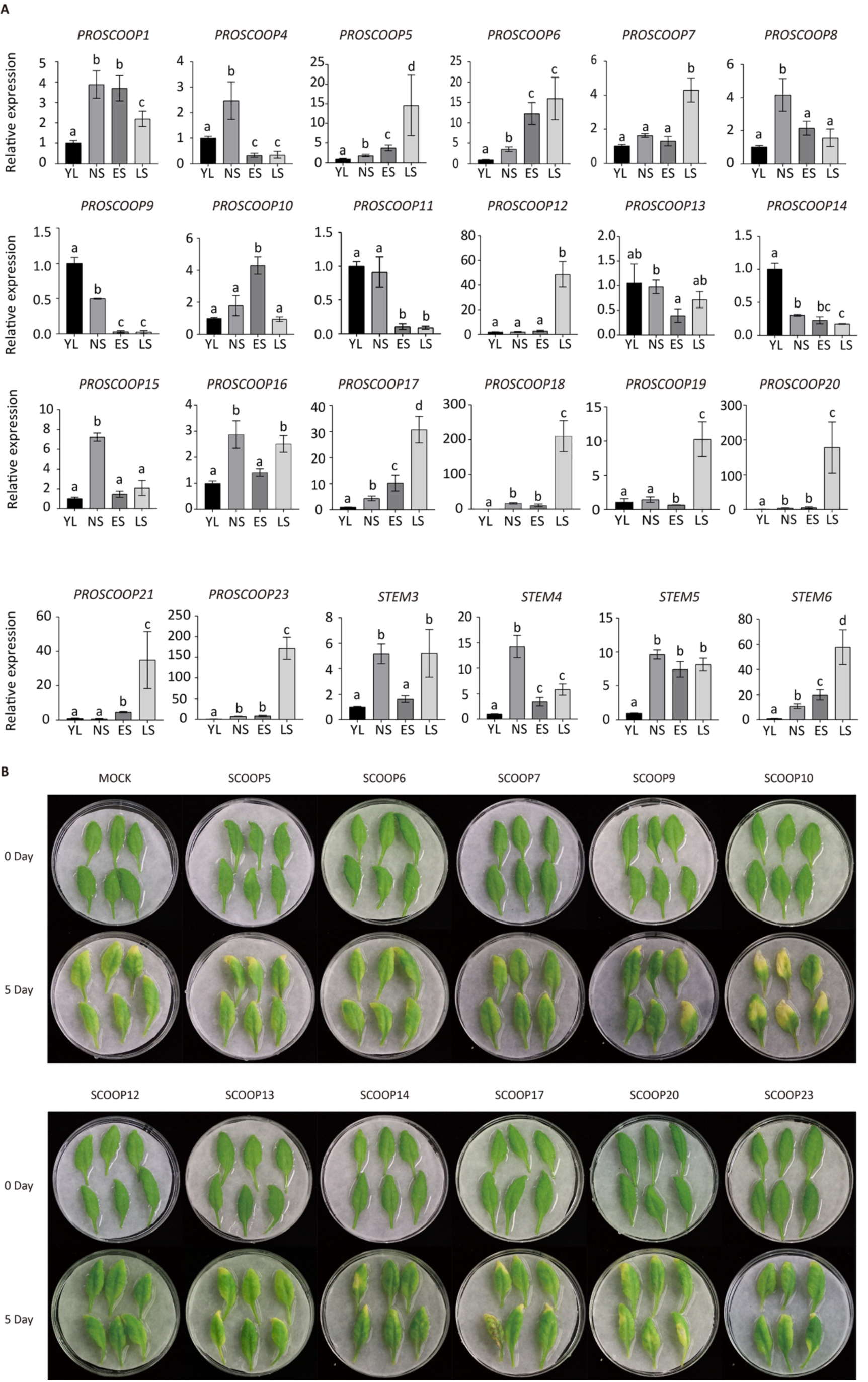
(A) Relative expression of *PROSCOOP*s at different leaf senescence stages. The bars represent the means ± SD of three biological replicates. Different letters represent statistically significant difference analyzed by one-way ANOVA analysis (P < 0.05). (B) Senescence phenotype analyzed in the 5^th^ and 6^th^ rosette leaves from 4-week-old WT plants upon exogenous application of synthetic SCOOP peptides (1 µM). Three replicates were performed with 6 leaves analyzed for each replicate.

**Figure S3.**
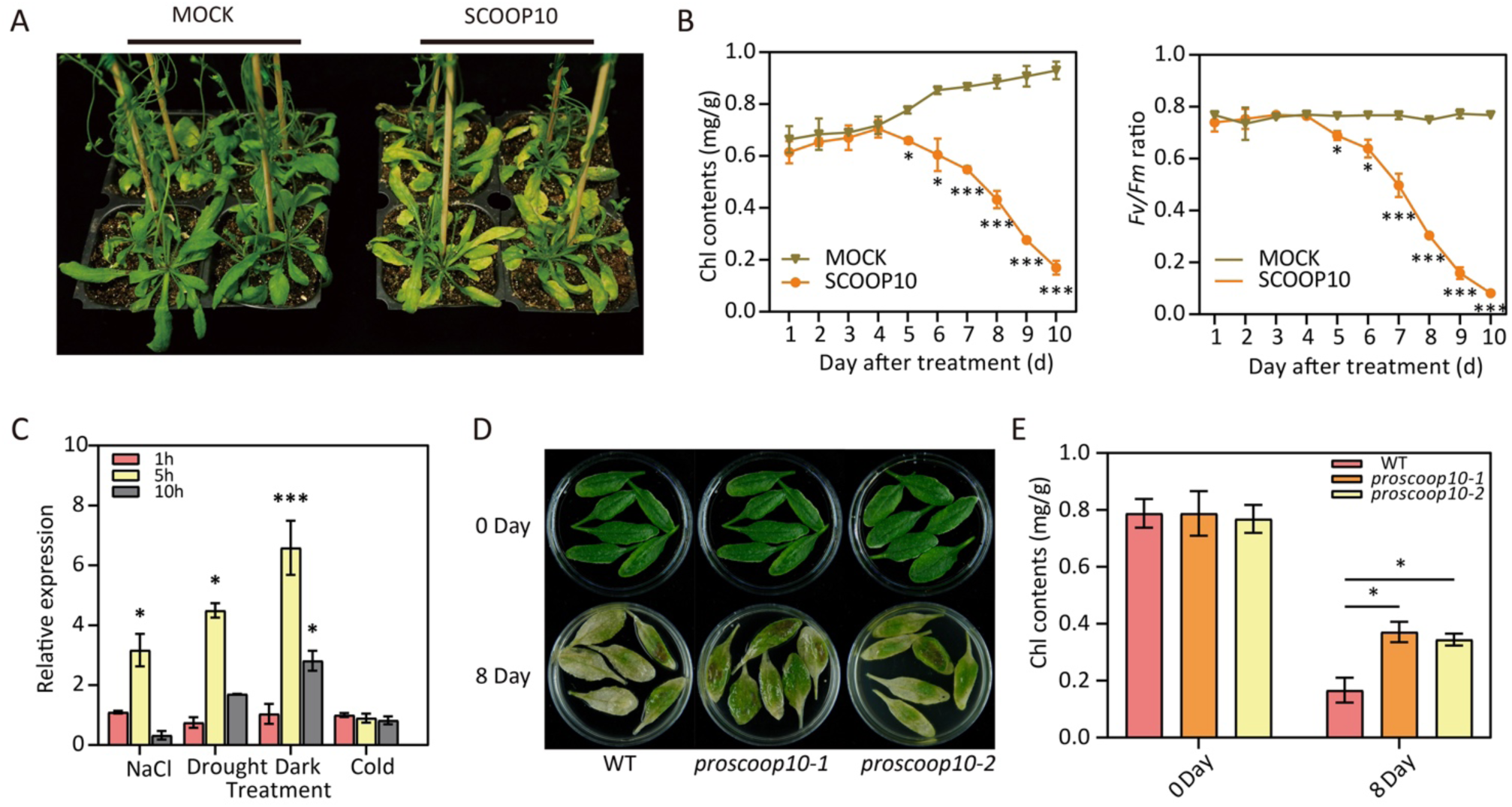
(A) Senescence phenotype of 4-week-old WT plants sprayed with synthetic SCOOP10 peptides (1µM) or ddH2O (MOCK) for 10 days. (B) Measurement of Chl content and ion leakage of the 6^th^ rosette leaves from plants shown in (A) after 1 to 10 days of treatments. Three biological replicates were performed with 8 leaves measured for each replicate. (C) Relative expression of *PROSCOOP10* analyzed in the 6^th^ rosette leaves detached from 4-week-old WT plants after treatments with diverse abiotic stresses (150 mM NaCl, 300 mM mannitol, darkness and 4°C) for 1, 5 and 10 hours. Three biological replicates were included for each experiment. (D) Senescence phenotype of the 6^th^ rosette leaves from 4-week-old WT plants incubated in ddH2O upon darkness treatments for 8 days. (E) Chl content measurement of detached leaves present in (D) at indicated time points. Three replicates each containing 6 leaves were analyzed. The error bars represent ± SD of 3 biological replicates with three technical replicates for each experiment. (*p < 0.05 and ***p < 0.01).

**Figure S4.**
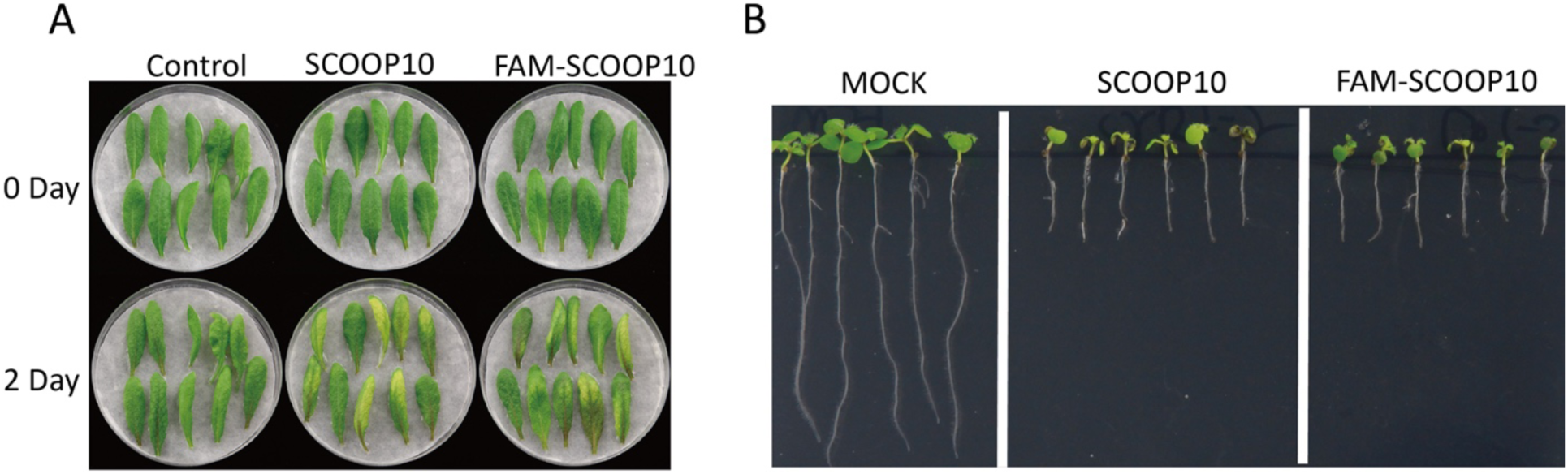
Identification of the bioactivity of fluorescently labeled FAM-SCOOP10 peptides. (A) Leaf senescence was analyzed in rosette leaves from 4-week-old plants treated with FAM- SCOOP10 and SCOOP10 peptides. (B) FAM-SCOOP10 induced root growth inhibition similarly to SCOOP10. 4-day old seedlings were transferred to 1/2 MS plates containing MOCK (ddH2O), 0.5 µM SCOOP10 or FAM- SCOOP10 peptides and grown for 7 additional days under long-day conditions.

